# Perforin-2 limits pathogen proliferation at the maternal-fetal interface

**DOI:** 10.1101/2020.05.20.107193

**Authors:** Petoria Gayle, Vanessa McGaughey, Rosmely Hernandez, Marina Wylie, Rachel C. Colletti, Ka Lam Nguyen, Marshall Arons, Laura Padula, Natasa Strbo, Kurt Schesser

## Abstract

Placental immune responses are highly regulated to strike a balance between protection and tolerance. For relatively mild infections, protection encompasses both the mother and fetus; however, during worsening conditions, protection becomes exclusively reserved for the mother. Previously, we and others have shown that the host factor Perforin-2 plays a central role in protecting mice and cells against infection. Here, we analyzed Perforin-2 activity in the mouse placenta to determine whether Perforin-2 plays a similarly protective role. We show that Perforin-2 is critical for inhibiting *Listeria monocytogenes* colonization of the placenta and fetus and that this protection is due to both maternal and fetal-encoded Perforin-2. *Perforin-2* mRNA is readily detectable in individual immune cells of the decidua and these levels are further enhanced specifically in decidual macrophages during high-dose infections that result in fetal expulsion. Unexpectedly, inductive Perforin-2 expression in decidual macrophages did not occur during milder infections in which fetal viability remained intact. This pattern of expression significantly differed from that observed in splenic macrophages in which inductive Perforin-2 expression was observed in both high and mild infection conditions. In the placenta, inductive Perforin-2 expression in decidual macrophages was co-incident with their polarization from a M2 to M1 phenotype that normally occurs in the placenta during high-burden infections. Our results suggest that Perforin-2 is part of a host response that is protective either for both the mother and fetus in milder infections or exclusively for the mother during high-dose infections.

## Introduction

The balance between host defense and tolerance during pregnancy is achieved both by the modulation of the maternal immune system towards the semi-allogenic fetus as well as through the barrier function of the placenta. Despite these safeguards, infections contribute approximately 25% of stillbirths in the United States, largely due to the ability of specific pathogens, including *Listeria monocytogenes*, to colonize the placenta (McClure et al. 2010). *L. monocytogenes* is widely used in hematogenous infection models to study host-pathogen interactions in the placenta (Lamond and Freitag 2018). While these models have advanced our understanding of the pathogenesis of placental infections, there still remains the need to further understand the mechanisms of immune defense in coordination with fetal tolerance during placental infection.

Numerous studies have shown that infections during pregnancy can disrupt the highly controlled inflammatory response and result in pregnancy complications such as miscarriages or spontaneous abortion (Kim et al. 2005; Mor et al. 2017; Rodrigues-Duarte et al. 2018). These pregnancy related complications are commonly attributed to the activation of the innate immune response. Specifically, macrophages of the decidua, the maternal component of the placenta, have been shown to be highly dynamic during pregnancy. These cells change and respond to the inflammatory environment of the placenta expressing characteristics of the classically activated phenotype, M1, in early and late pregnancy and resembling the M2 phenotype during the mid-stage of pregnancy (Shapouri-Moghaddam et al. 2018; Zhang et al. 2017). Excessive levels of pro-inflammatory M1 macrophages have been linked to abnormal pregnancy outcomes including pre-term labor and fetal mortality (Jena et al. 2019; Svensson-Arvelund and Ernerudh 2015; Wang et al. 2011; Xu et al. 2016). It has also been shown that *L. monocytogenes* can trigger M1 polarization in the placenta (Benoit et al. 2008). Studying the role of innate immune factors within the placenta is important to enhance our understanding of how to prevent the devastating effects of infection during pregnancy.

In vertebrates there are three pore-forming factors that protect against microbial pathogens. Complement component 9 (C9) and Perforin-1 possess membrane-attack-complex-perforin (MACPF) domains that mediate their polymerization and pore formation. The third and most recently identified vertebrate MACPF-containing factor, Perforin-2, is found in the earliest evolved animals, and is ancestral to C9 and Perforin-1 (D’Angelo et al. 2012). Unlike the Complement Factors and Perforin-1 that are secreted from cells, Perforin-2 is an integral membrane protein. In human and mouse cells *Perforin-2* mRNA is constitutively present in macrophages and is induced in fibroblastic cells following infection or exposure to inflammatory signals (McCormack et al. 2013). Perforin-2 plays a major role in protecting mice from *Listeria, Salmonella, Staphylococcus*, and *Yersinia* infections (McCormack et al. 2015a; 2015b; 2016). *In vitro*, Perforin-2 restricts intracellular *L. monocytogenes* proliferation by a pH-dependent mechanism in both primary peritoneal macrophages and fibroblastic cells suggesting that the rapid development of listeriosis in *Perforin-2* -/- mice is due to defects in cellular killing activity (McCormack et al. 2016). Recently it has been shown that Perforin-2 directly impacts type I interferon signaling by physically interacting with the IFN-α and –β receptors 1 and 2 (McCormack et al. 2020). Whether the bactericidal activity of Perforin-2 involves the interferon signaling machinery remains to be determined.

Here we tested whether Perforin-2 plays a protective role in limiting colonization of *L. monocytogenes* in the mouse placenta. Additionally, we analyzed *Perforin-2* mRNA expression in individual cells of the placenta following infection to determine whether *Perforin-2* expression levels change during infection.

## Materials and Methods

### Mice, microbes, and infections

Wild-type BALB/c and C57BL/6 *Perforin-2* +/+ and *Perforin-2* -/- littermates (McCormack et al. 2015a) were bred in the animal care facility at the University of Miami Miller School of Medicine, Miami, FL. All animal procedures were approved by the Institutional Animal Care and Use Committee, University of Miami Miller School of Medicine (protocol 19-075). Animals were housed under a circadian cycle (12hr light/12hr dark cycle). Virgin female mice were mated between 6 - 12 weeks of age, and checked daily for estrous stage and copulatory plugs. Presence of a plug was denoted as gestation day (GD) 0.5. Weight gain was monitored on GD 4.5 through GD 12.5 to confirm pregnancy. *L. monocytogenes* (10403S) were grown in brain heart infusion media at 37 °C with vigorous shaking to mid-log phase. Mice at GD 12.5 were infected intravenously with doses as indicated in figure legends. At 44 hours post infection (hpi), mice were humanely euthanized and uterine horns, livers, and spleens were removed and processed for either single cell analysis (see below) or colony forming unit (CFU) assays. For CFU assays, placentas (including decidual tissue), fetuses, and livers were further dissected and homogenized using a fine wire mesh to grind the tissues in sterile water containing 0.05% Tween. The resulting tissue homogenates were diluted and plated on Luria Broth agar to determine *L. monocytogenes* titers. In the experiments using heterogenic matings, fetal tissue homogenates were used for genotyping.

### Preparation of single-cell suspensions

Following their removal, uterine horns were dissected to remove individual fetal-placental units (FPU), each FPU was further dissected to isolate decidual tissue. Livers (minced) and pooled deciduae were incubated in 2 mg/mL collagenase D (Roche) at 37 °C for 30 – 40 minutes with agitation. The resulting cell suspensions were then washed with cold IMDM (Life Technologies) + 10% heat-inactivated FBS (Sigma-Aldrich), passed through a 70 μm filter followed by passaging through a 40 μm filter (VWR International, Radnor, PA). Spleens were gently homogenized through 70 μm filters and similarly washed and passed through 40 μm filters. The resulting single cells were then centrifuged at 500g for 5 min at 4°C, treated with ACK Lysing Buffer (Life Technologies Corporation) for 5 minutes to remove red blood cells, and finally washed with IMDM + 10% FBS. Total viable cells were determined using the Vi-Cell XR Cell Viability Analyzer (Beckman Coulter).

### Single cell analysis

Isolated cells were washed in FACS staining buffer and incubated with anti-CD16/32 (clone 2.4G2) to block FcRs for 10 mins followed by an incubation with Live/Dead fixable yellow dead cell stain (ThermoFisher) and fluorescent-conjugated monoclonal antibodies (mAbs) for 30 min. The following anti-mouse mAbs were used for analysis: CD45 (30-F11; BioLegend), CD3 (17A2; BioLegend), F4/80 (BM8; BioLegend), CD11b (M1/70; BioLegend), Ly6C (HK1.4; BioLegend), Ly6G (1A8; BioLegend), CD335 (29A1.4; BioLegend), MHC II (M5/114.15.2; BD Biosciences), CD206 (C068C2; BioLegend). Depending on the experiment, samples were either immediately analyzed by flow cytometry using a Sony SP6800 Spectral Analyzer or further processed to determine *Perforin-2* mRNA levels. Branched oligonucleotide signal amplification was used to determine *Perforin-2* mRNA levels in individual cells (PrimeFlow; ThermoFisher Scientific). Briefly, single cell suspensions were stained for surface antigens as described above, then fixed, permeabilized, and incubated with probes specific for *Perforin-2* transcripts (Assay Id: VB1-20172-PF; ThermoFisher Scientific). Cells were then subjected to a series of signal amplification cycles and then analyzed by flow cytometry as described above using FlowJo software (BD Biosciences).

## Results

### Perforin-2 limits pathogen colonization at the maternal-fetal interface

Pregnant wild-type and isogenic Perforin-2-deficient (*Perforin-2* -/-) mice were infected intravenously on gestation day (GD) 12.5 (i.e., mid-gestation) with *L. monocytogenes*. At 44 hours post infection, dams were humanely euthanized and livers and fetal placental units (FPU) were evaluated for *L. monocytogenes* by colony forming unit (CFU) assay. In infected *Perforin-2* +/+ dams, 54% (12/22) of the placentas and 14% (3/22) of fetuses possessed detectable levels of *L. monocytogenes* (**Fig. 1**). Dosages that result in approximately 50% of placentas in *Perforin-2* +/+ mice becoming infected will henceforth be referred to as placental dose 50 (PD_50_). In contrast, *Perforin-2* -/- mice harboring comparable levels of *L. monocytogenes* in the liver as *Perforin-2* +/+ dams, 88% (15/17) of the placentas and 71% (12/17) of fetuses possessed detectable levels of *L. monocytogenes*. These results suggest that Perforin-2 may play a significant role in limiting *L. monocytogenes* colonization of the placenta and fetus.

**Fig. 1.**
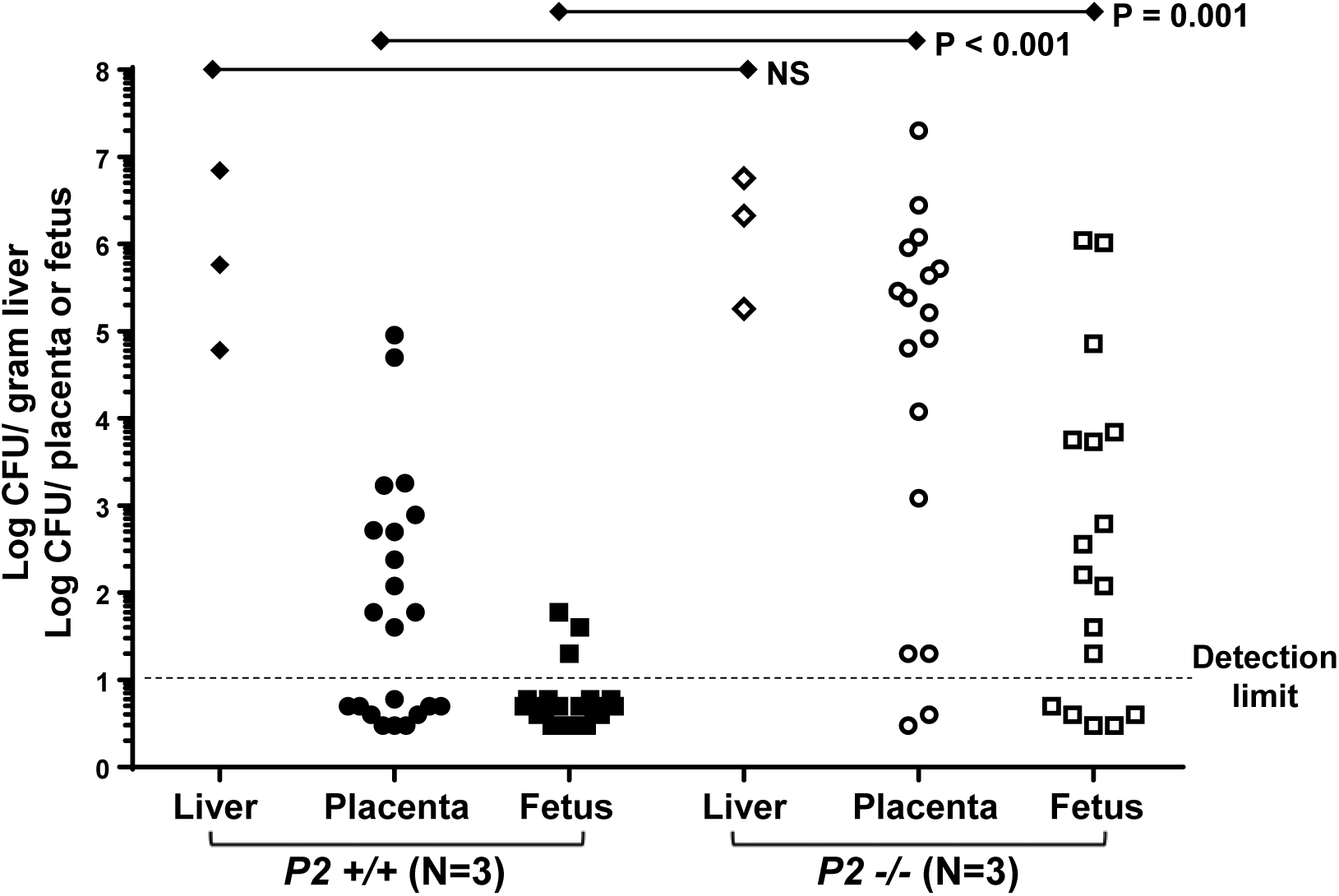
Perforin-2 limits *L. monocytogenes* infection of the placenta and fetus. Pregnant *Perforin-2* (*P2*) *+/+* and *-/-* mice were infected intravenously on GD 12.5 with 2.5 × 10^5^ CFU of *L. monocytogenes* for 44 hours. Bacterial loads were then determined in each individual liver, placenta, and fetus by CFU assay. Shown are the compiled results of two separate infection experiments using a total of 3 pregnant mice per genotype harboring a total of either 22 FPUs (*P2* +/+) or 17 FPUs (*P2* -/-). Mann-Whitney U test used to calculate p-values.

### Maternal and fetal-derived Perforin-2 contributes to protection against infection in the placenta

The placenta is a chimeric organ that consists of maternally-derived tissue and fetal-derived trophoblast cells. A heterogenic mating strategy was used to evaluate the specific contributions of maternal- and fetal-derived Perforin-2 in limiting *L. monocytogenes* colonization in the placenta. Initially, *Perforin-2 -/-* female mice were crossed to *Perforin-2 +/-* males (generating approximately 50% *Perforin-2 +/-* and 50% *Perforin-2 -/-* fetuses) and at GD 12.5, dams were infected with PD_50_ *L. monocytogenes* and analyzed as described above. A representative dam is shown in which the infection burdens of FPUs containing *Perforin-2* +/- fetuses are generally lower than FPUs containing *Perforin-2* -/- fetuses (**Fig. 2A**). In compiled data from 5 dams, the placentas associated with *Perforin-2* +/- fetuses (designated as T(+/-)) had significantly lower infection burdens compared to placentas associated with *Perforin-2* -/- fetuses (designated as T(-/-)). A similar reduction in infection burden was observed in *Perforin-2* +/- fetuses compared to *Perforin-2* -/- fetuses (**Fig. 2B**) indicating that fetal-derived Perforin-2 protects the placenta from infection. In reciprocal matings, in which *Perforin-2 +/-* female mice were crossed with *Perforin-2 -/-* males and infected at GD 12.5, a more modest protective effect of fetal-derived Perforin-2 was observed (**Fig. 2C**), possibly indicating that maternal-derived Perforin-2 partially masks the protective effect of fetal-derived Perforin-2. Collectively these data show that both maternal- and fetal-derived Perforin-2 contribute to protecting the placenta from being colonized by a bloodborne pathogen.

**Fig. 2:**
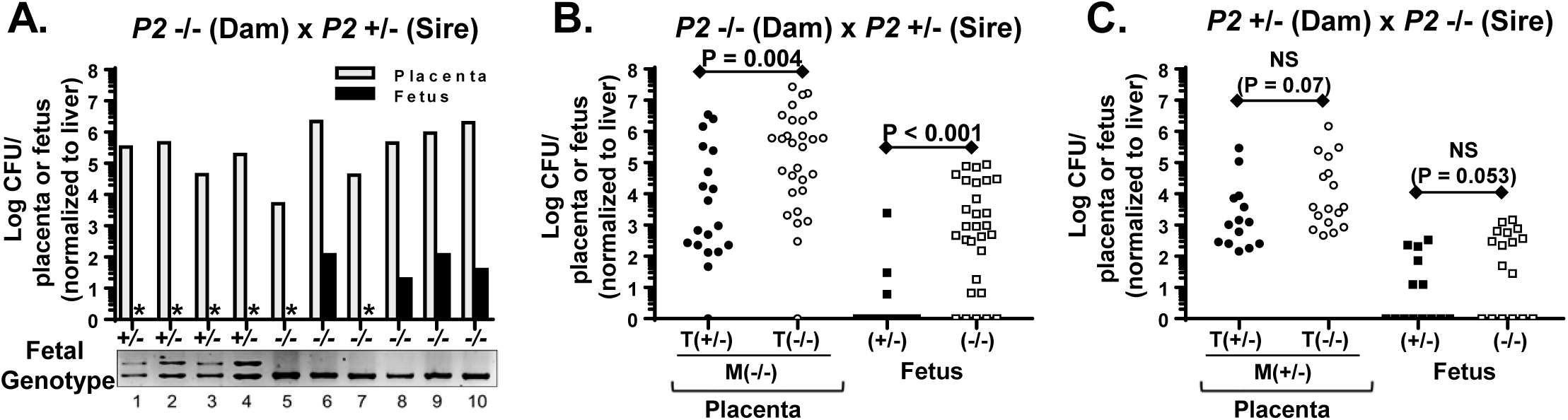
Both maternal- and fetal-encoded Perforin-2 contribute to protection of the placenta and fetus. A heterogenic mating strategy was used to generate pregnant mice that possessed fetuses that were either *Perforin-2* (*P2*) *+/-* or *-/-*. Pregnant mice (GD 12.5) were infected intravenously with 2.5 × 10^5^ CFU *L. monocytogenes* for 44 hrs. **(A)** A representative cross is shown between a *P2 -/-* female and a *P2 +/-* male. Following infection, FPUs (N=10) were dissected, bacterial loads determined in each individual placenta and fetus by CFU assay, and individual fetuses genotyped. In the example shown, fetuses #1-4 are *P2 +/-* and #5-10 are *P2 -/-*. Asterisk denotes CFU values below detection limit. **(B)** Compiled analysis of crosses between *P2 -/-* females, N=5 and *P2 +/-* males in which FPUs lack maternal-derived P2, designated as M(-/-). Placentas associated with *P2 +/-* fetuses (e.g., fetuses 1-4 in **(A)**) possess trophoblasts T(+/-) with fetal-derived P2. Placentas associated with *P2 -/-* fetuses (e.g., fetuses 5-10 in **(A)**) possess trophoblasts T(-/-) that lack fetal-derived P2. **(C)** Similar compiled analysis of crosses between *P2 +/-* females, N=5 and *P2 -/-* males in which FPUs possess maternal-derived P2, designated as M(+/-). Placentas associated with *P2 +/-* fetuses possess trophoblasts, designated as T(+/-), with fetal-derived P2. Placentas associated with *P2 -/-* fetuses possess trophoblasts, designated as T(-/-), that lack fetal-derived P2. Compiled data drawn from three independent infections performed on separate days. Mann-Whitney U test was used to calculate p-values. (NS = not significant)

### Perforin-2 expression is induced in placental immune cells following infection

The maternal component of the placenta, the decidua, is the initial colonization site of various bloodborne pathogens including *L. monocytogenes* (Rizzuto et al. 2017). We initially analyzed the cellular composition of the decidua and the liver in uninfected and infected mid-gestation mice. Previously we and others showed that the immune cell composition in the liver in non-pregnant mice undergoes substantial changes following *L. monocytogenes* infection (Gregory et al. 2002; Blériot et al. 2015; Gayle et al. 2019). There was a similar pattern of changes in the liver observed in mid-gestation pregnant mice following infection, including a marked disappearance of resident macrophages (CD11b^lo^/F4/80^+^) and the infiltration of inflammatory monocytes and neutrophils (CD11b^+^/F4/80^-^) (**Fig. 3A**). In contrast, in the decidua of the same mice, there were no significant changes in the resident macrophages (CD11b^+^/F4/80^+^) or other cell types following infection (**Fig. 3B**). From GD 12.5 – 14.5 pregnant mice approximately a million cells are typically isolated from individual decidua of which ∼20% are CD45^+^. The 4 × 10^5^ CD45^+^cells per decidua is composed of approximately 20% CD11b^-^/F480^-^ (primarily NK cells), 30% CD11b^+^/F480^-^ (monocytes and neutrophils), and 40% CD11b^+^/F480^+^ (macrophages).

**Fig. 3:**
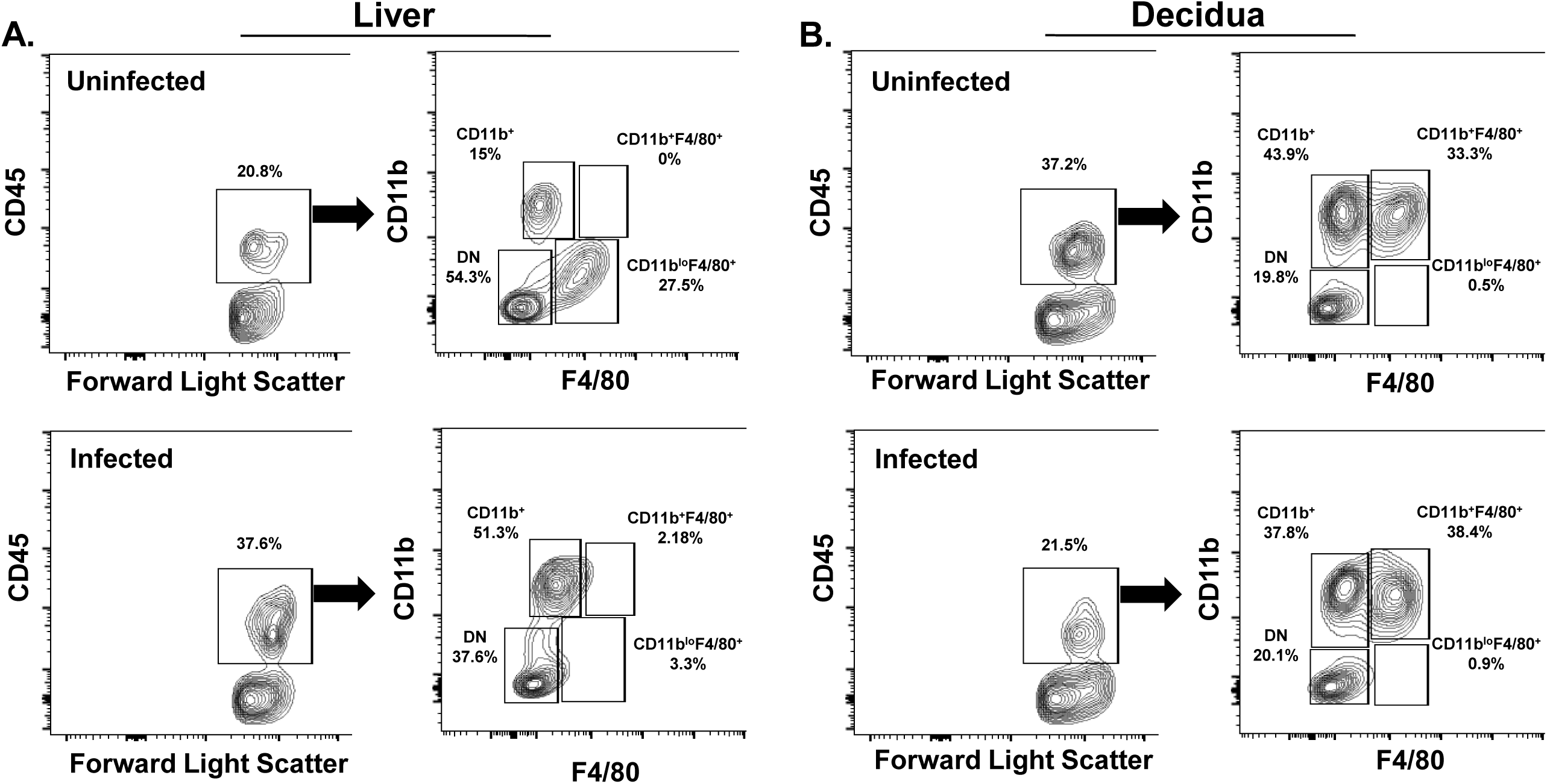
Composition of immune cells in liver and decidua of uninfected and infected GD12.5 pregnant mice. Pregnant mice (GD 12.5) were either uninfected or intravenously infected with 1 × 10^6^ CFU of *L. monocytogenes* for 44 h. Single-cell preparations of the indicated tissue were analyzed by flow cytometry for expression of immune- and myeloid-specific markers. **(A)** The percentage of cells staining positive for the immune-specific cell surface marker CD45 per 10^6^ isolated total live liver cells shown from an uninfected and infected mouse. The left panel shows the staining profile of CD45-staining liver cells from an individual uninfected dam (top) and an individual infected dam (bottom). The right panel shows the expression levels of myeloid-specific markers CD11b and F4/80 of the CD45^+^ cells. **(B)** For the same mice, the percentage of cells staining positive for the immune-specific cell surface marker CD45 per 10^6^ isolated total live decidual cells are shown. The left panel shows the staining profile of CD45-staining decidual cells from an individual uninfected dam (top) and an individual infected dam (bottom) and the right panel shows the expression levels of myeloid-specific markers CD11b and F4/80 of the CD45^+^ cells. Shown is a representative mouse from 3 mice per group.

*Perforin-2* mRNA levels were assayed in individual decidual cells to determine both cell type-specific expression and whether expression levels change following infection. To ensure that all placentas within each pregnant mouse became colonized by *L. monocytogenes* during the 44 hr infection period, the doses used for these experiments will be referred to as placental dose 100% (PD_100_) that are 2- to 4-fold higher than the ‘low-dose’ experiments shown in Figs. 1 and 2. As described earlier, pregnant mice (GD 12.5) were either left uninfected or infected with *L. monocytogenes* and following 44 hours of infection, FPUs were collected and placentas were further dissected to isolate decidual cells. Pooled decidual cells from each individual dam were analyzed for cell surface markers and *Perforin-2* mRNA levels. In uninfected dams, *Perforin-2* mRNA was readily detected in CD45^+^ decidual cells and this signal was enhanced 2- to 3-fold in CD45^+^ decidual cells isolated from infected dams (**Fig. 4A, B**). A similar 2- to 3-fold infection-dependent increase in *Perforin-2* mRNA levels was also observed in CD45^+^ splenic cells derived from the same mice (**Fig. 4C**).

**Fig. 4:**
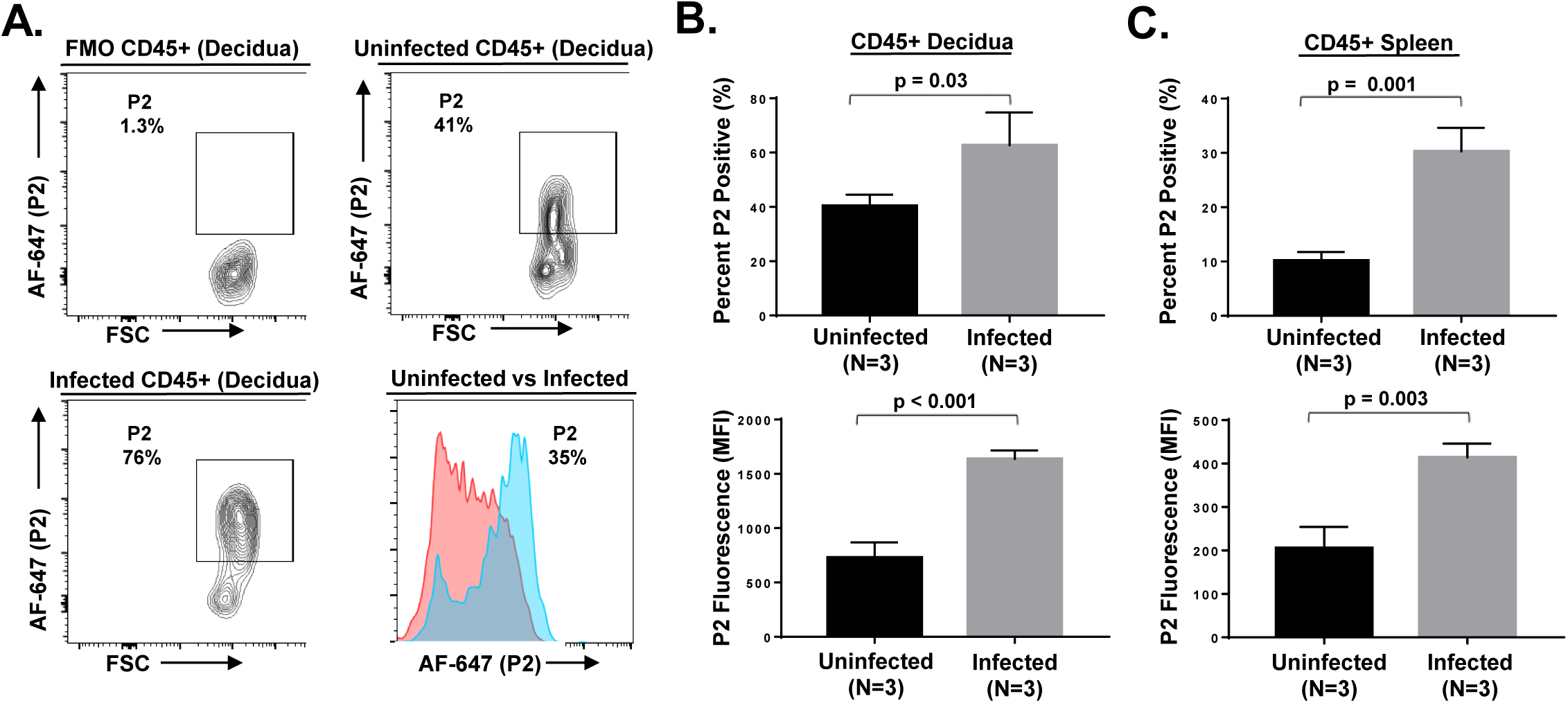
*Perforin-2* expression is induced in decidual and splenic immune cells following infection. Pregnant BALB/c mice (GD12.5) were infected intravenously with 5 × 10^5^ - 1 × 10^6^ CFU of *L. monocytogenes* for 44 hrs and decidual and splenic cells were analyzed for *P2* mRNA levels. **(A)** Representative contour plots of background (FMO), uninfected, and infected cells gated on live, CD45^+^ cells expressing AF-647 (P2). Histogram overlay of uninfected (red) and infected (blue) CD45^+^ decidual cells showing *Perforin-2* (*P2*) mRNA levels. **(B)** Compiled analysis of *P2* percentage and mean fluorescence intensity (MFI) of *P2* expression on CD45^+^ cells in decidua and spleen. Compiled data drawn from two independent infections performed on separate days. Student’s T-test was used to calculate p-values.

Of the three major CD45^+^ subsets found in the GD 12.5 decidua of BALB/c mice (see **Fig. 3B**), infection-induced enhancement of *Perforin-2* mRNA levels occurred primarily in the CD11b^+^F4/80^+^ macrophages (**Fig. 5A**,**B**). This infection-induced increase also occurred in the CD45^+^ splenic macrophages derived from the same mouse (**Fig. 5C**). In addition, *Perforin-2* mRNA was also specifically induced in infected decidual and splenic CD11b^+^F4/80^+^ macrophages isolated from C57BL/6 mice (**Fig. 5D-F**). These data show that *Perforin-2* mRNA levels increase in immune cells of the decidua following infection and that this increase primarily occurs in macrophages.

**Fig. 5:**
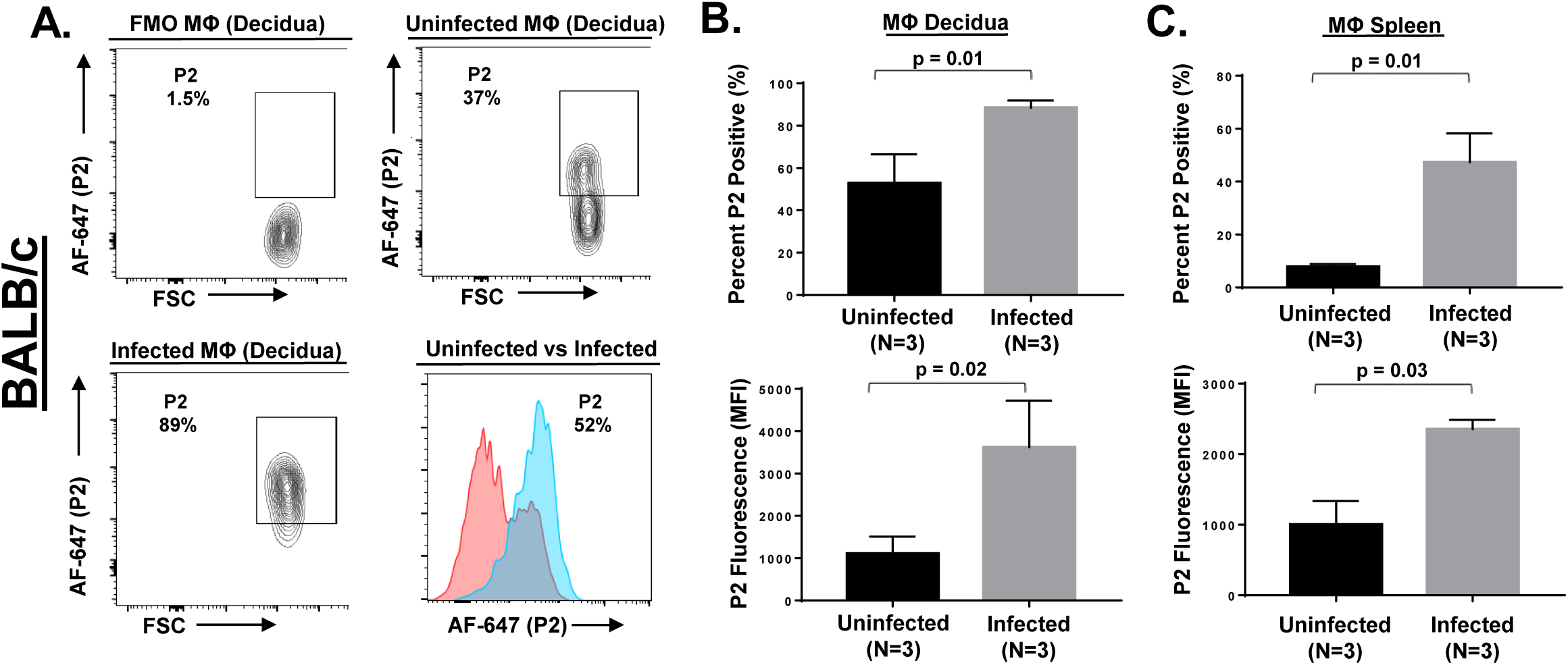
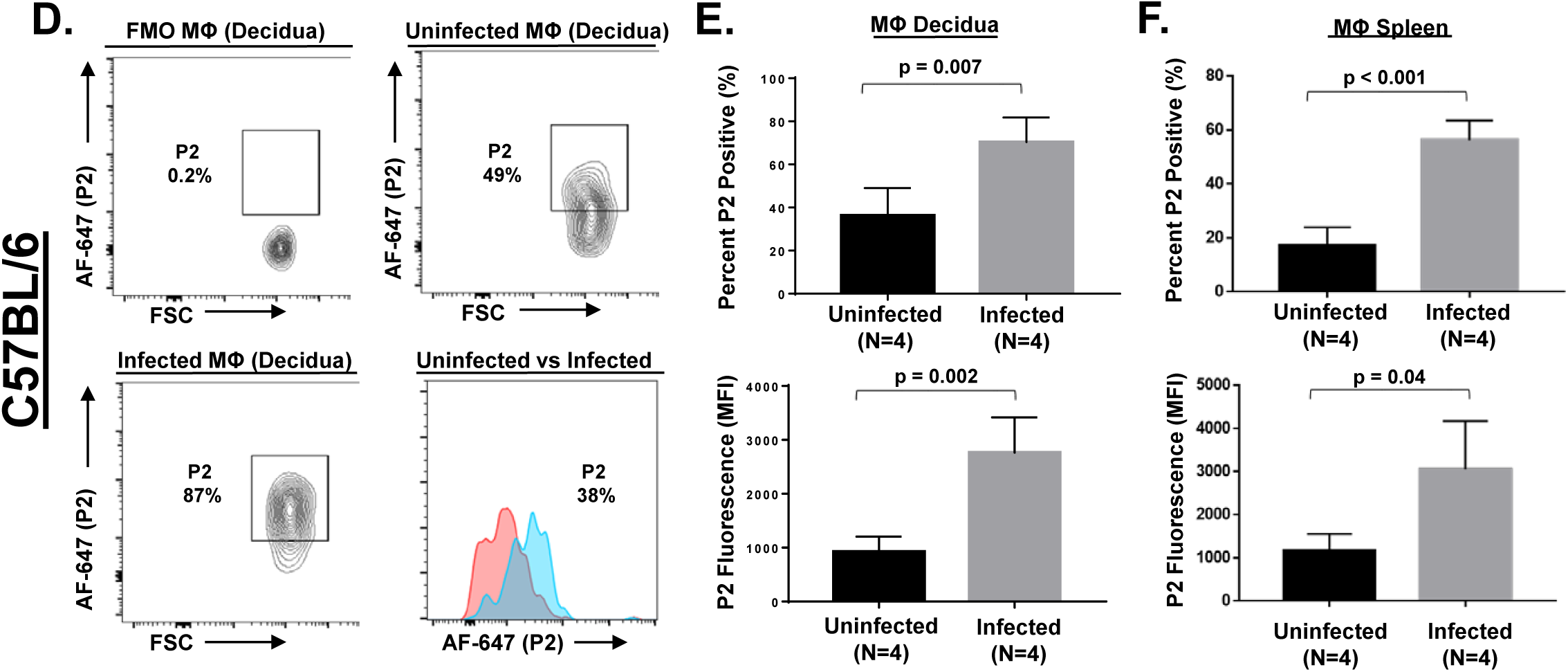
Specific *Perforin-2* induction in decidual and splenic macrophages of BALB/c and C57BL/6 dams following infection. Pregnant BALB/c (**A-C**) and C57BL/6 (**D-F**) mice (GD12.5) were infected intravenously with 5 × 10^5^ - 1 × 10^6^ CFU of *L. monocytogenes* for 44 hrs and decidual and splenic macrophages were analyzed for *Perforin-2* (*P2*) mRNA levels. **(A**,**D)** Representative contour plots of background (FMO), uninfected, and infected cells gated on live macrophages expressing AF-647 (P2). Histogram overlay of uninfected (red) and infected (blue) CD45^+^ decidual macrophages showing *P2* mRNA levels. Compiled analysis of *P2* percentage and mean fluorescence intensity (MFI) of *P2* expression in decidual macrophages **(B**,**E)** and spleen **(C**,**F)**. Compiled data drawn from two independent infections performed on separate days. Student’s T-test used to calculate p-values.

### Reduced infection dosages result in modest changes in Perforin-2 mRNA levels in decidual macrophages

As noted earlier, the *Perforin-2* expression experiments shown in Figs. 4 and 5 used infective doses (PD_100_) that resulted in all placentas becoming colonized with *L. monocytogenes* by 44 hpi. Unexpectedly, when these experiments were performed with 2- to 4-fold lower doses of *L. monocytogenes* in which approximately 50% of placentas had become colonized by *L. monocytogenes* by 44 hpi (PD_50_), there was little to no difference in *Perforin-2* mRNA levels between decidual macrophages isolated from uninfected and infected GD 12.5 pregnant mice (**Fig. 6A**,**B**). In contrast, in the same mice *Perforin-2* mRNA levels were significantly elevated in splenic macrophages isolated from infected mice compared to uninfected mice (**Fig. 6C**). A similar pattern of *Perforin-2* expression in decidual and splenic macrophages was observed in pregnant GD 12.5 C57BL/6 mice infected at PD_50_ (**Fig. 6D**,**E**). These findings suggest that in the placenta there is a bacterial dose-dependent induction of *Perforin-2* mRNA.

**Fig. 6:**
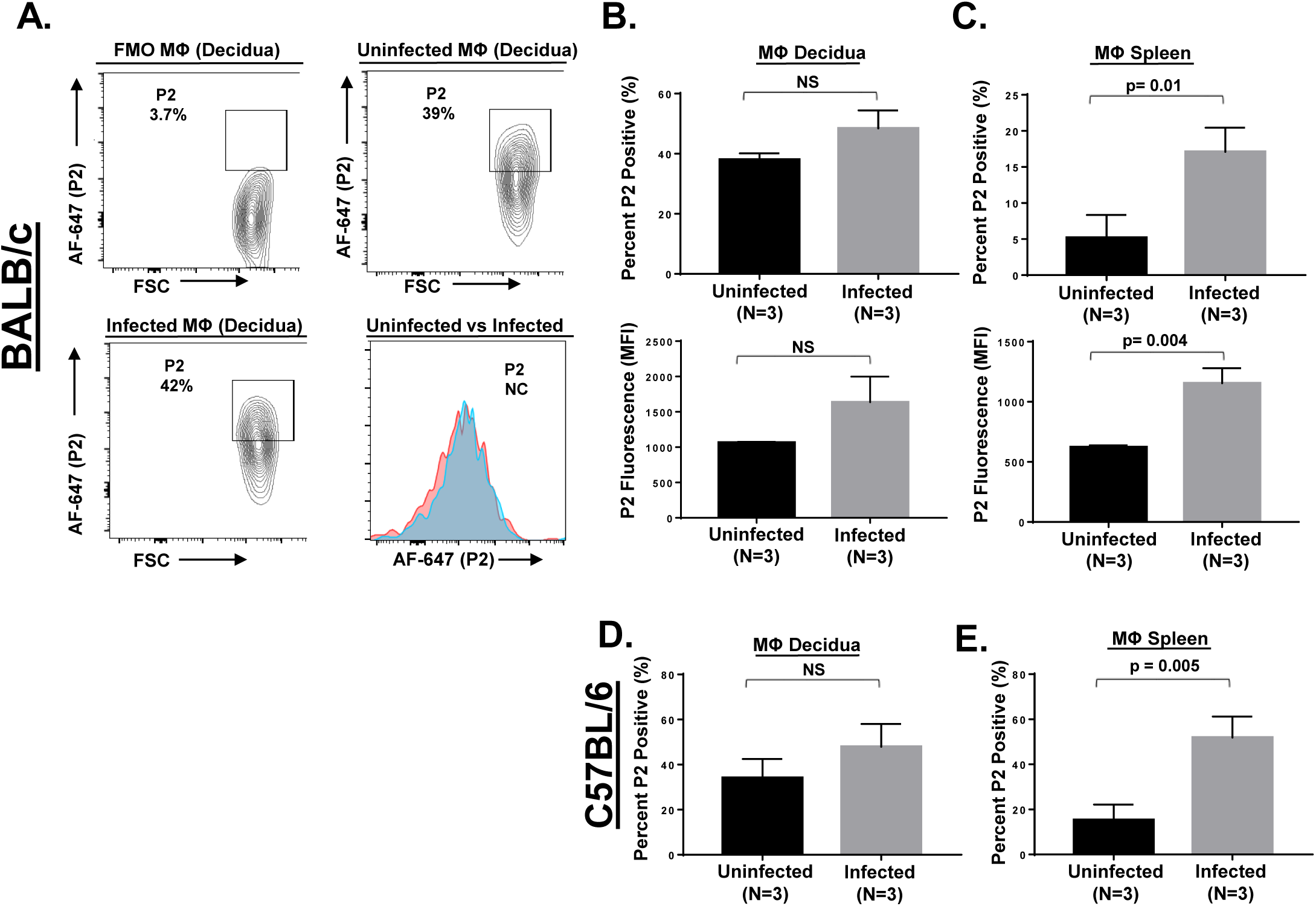
Divergent *Perforin-2* expression in decidual and splenic macrophages following lower-dosed infections. Pregnant BALB/c **(A-C)** or C57BL/6 **(D**,**E)** mice (GD 12.5) were infected intravenously with 2.5 × 10^5^ CFU of *L. monocytogenes* for 44 hrs and single-cell preparations of decidual and splenic macrophages were analyzed for *Perforin-2* (*P2*) mRNA levels and shown as described in Fig. 5. Student’s T-test used to calculate significance. (NS = not significant)

### Perforin-2 mRNA levels are elevated in M1 decidual macrophages

In humans and mice, decidual macrophages at mid-gestation are primarily of the M2 phenotype and polarization of these macrophages to the inflammatory M1 phenotype is associated with a variety of complications that can lead to premature pregnancy termination (Brown et al. 2014; Jena et al. 2019; Svensson-Arvelund and Ernerudh 2015; Xu et al. 2016). To determine the infection-specific phenotype of decidual macrophages in our model, pregnant GD 12.5 mice were either left uninfected or infected with *L. monocytogenes* at PD_100_. Following their isolation, decidual cells were stained for M1 and M2 specific markers and analyzed by flow cytometry. Macrophages were defined as single, live, CD45^+^, CD11b^+^, F4/80^+^, Ly6C^lo^, and Ly6G^-^ cells (**Fig. 7A)**. Macrophages were then further classified as either M1 (CD206^-^ MHCII^hi^) or M2 (CD206^+^ MHCII^lo^) in uninfected and infected dams. Consistent with previously published findings cited above, decidual macrophages isolated from uninfected dams were primarily M2. In contrast, decidual macrophages isolated from infected dams contained lower frequencies of CD206^+^ cells and higher frequency of MHCII cells indicative of a M1-skewed phenotype (**Fig. 7**). When GD 12.5 mice were infected with a reduced dose (PD_50_) of *L. monocytogenes*, the M1/M2 distribution of decidual macrophages at 44 hpi did not appreciatively differ from that observed in decidual macrophages isolated from uninfected GD 12.5 mice (*not shown*).

**Fig. 7:**
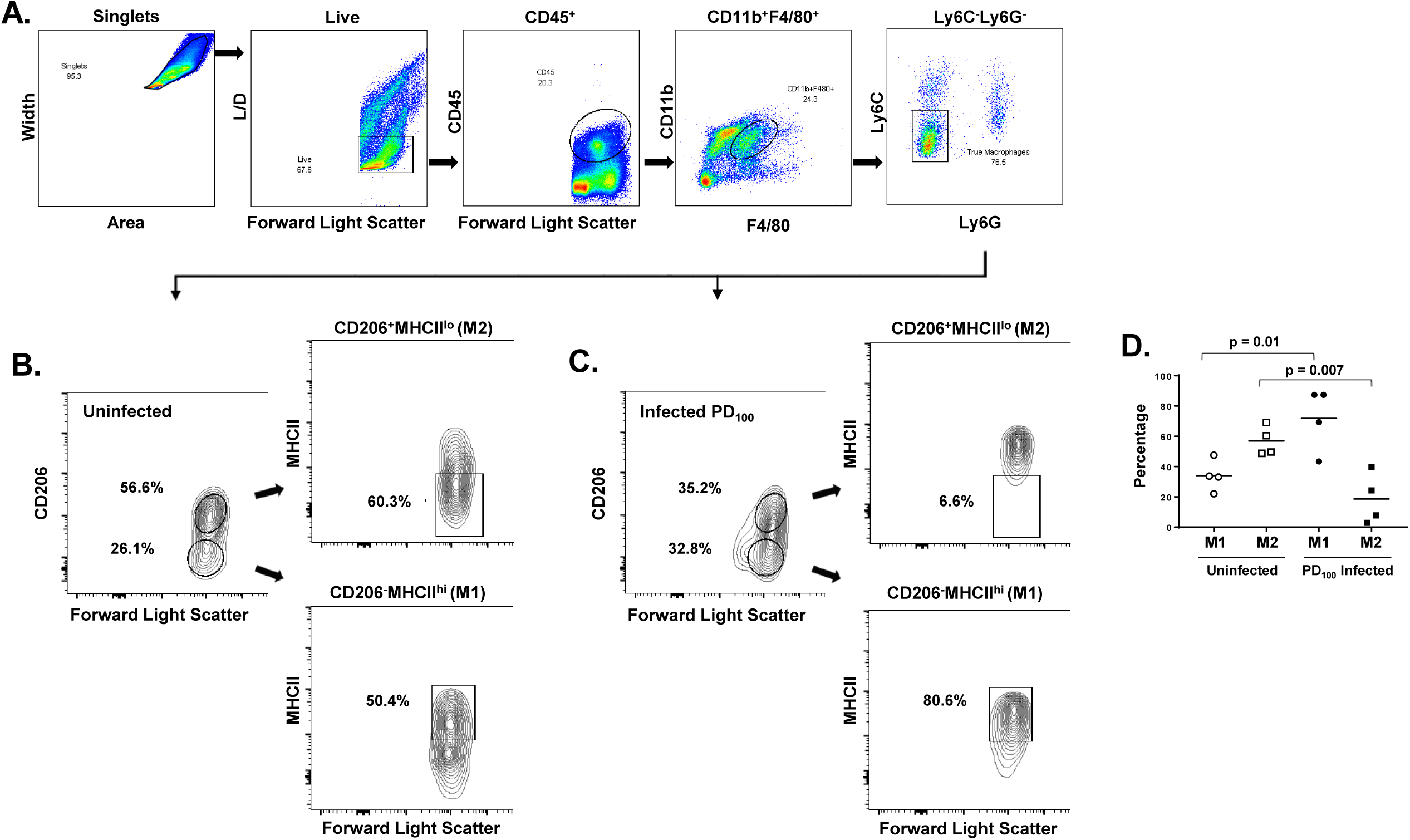
Decidual macrophages polarize to a M1 phenotype following high-dose infection. Pregnant mice (GD 12.5) were either uninfected or intravenously infected with 1 × 10^6^ CFU (PD_100_) of *L. monocytogenes* for 44 h. Single-cell preparations of deciduas were analyzed by flow cytometry for expression of M1 and M2 macrophage-specific markers. **(A)** Gating strategy used to define macrophages (single, live, CD45^+^, CD11b^+^, F4/80^+^, Ly6C^-^, Ly6G^-^). Gating strategy and proportions of M2 (CD206^+^, MHCII^lo^) and M1 (CD206^-^, MHCII^hi^) macrophages in a representative uninfected **(B)** and PD_100_ infected **(C)** dam (GD 12.5). **(D)** Compiled analysis of M1 and M2 decidual macrophages by percentage in uninfected and PD_100_ infected dams. Results based on 4 mice analyzed on several different days. Student’s T-test was used to calculate p-values.

Decidual macrophages isolated from uninfected dams at mid-term pregnancy (GD 12.5) were stained for M1 and M2 specific markers as described above and subsequently analyzed for *Perforin-2* mRNA levels. In every pregnant mouse examined, *Perforin-2* mRNA levels were notably elevated in M1 decidual macrophages compared to M2 decidual macrophages (**Fig. 8**). Collectively, these data show that at high pathogen burdens M1 macrophages with heightened *Perforin-2* mRNA levels predominate in the placenta.

**Fig. 8:**
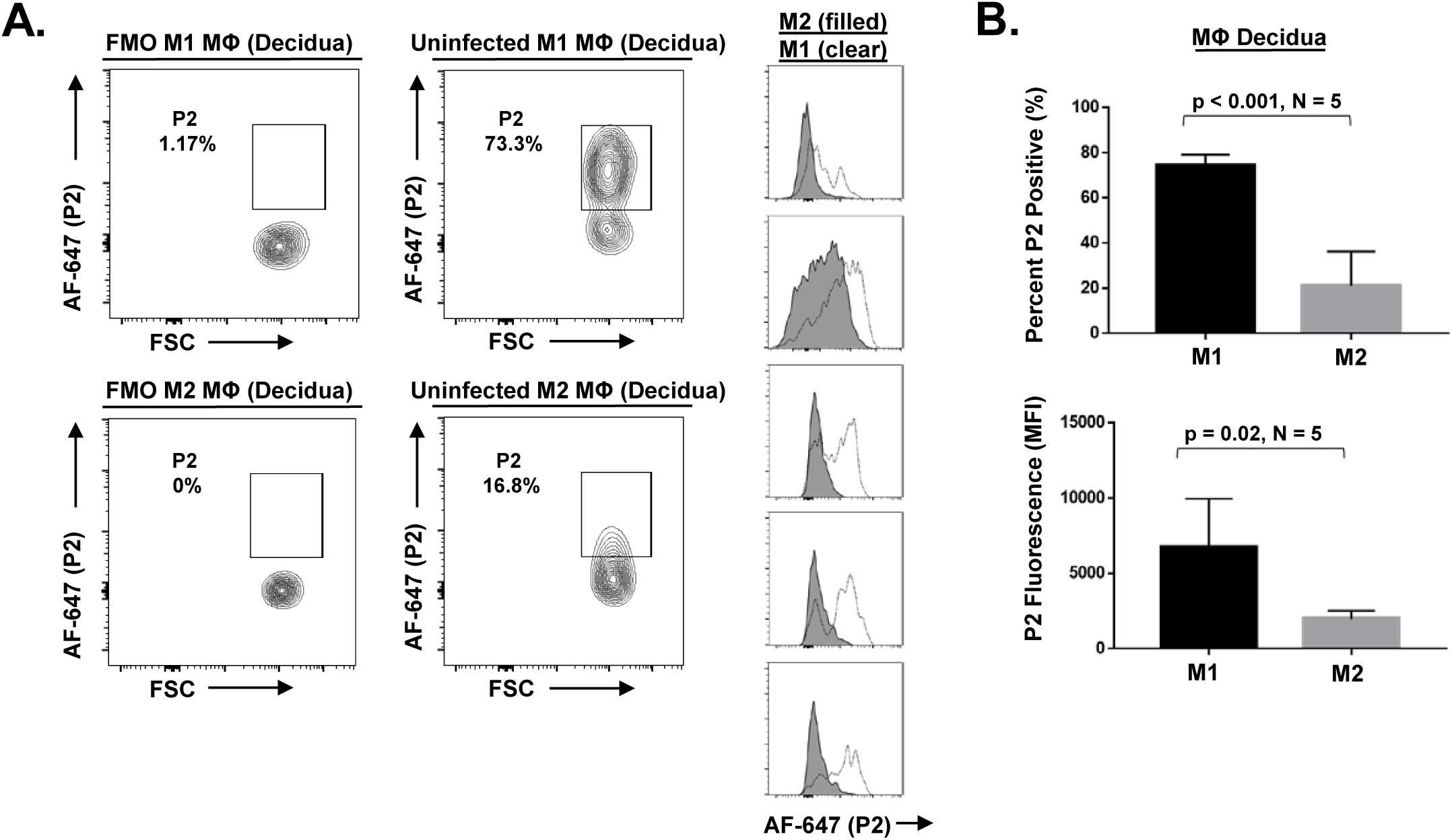
*Perforin-2* is expressed preferentially in decidual M1 macrophages. Decidual cells were isolated from uninfected pregnant *Perforin-2* (*P2*) *+/+* BALB/c mice (GD 12.5) and M1 and M2 macrophages were analyzed for *P2* mRNA. **(A)** Representative contour plots are shown of background (FMO) and uninfected M1 (top) and M2 (bottom) macrophages expressing AF-647 (P2). Histogram overlay plots of individual dams showing *P2* mRNA levels in M1 (clear) and M2 (grey filled) decidual macrophages. **(B)** Compiled analysis of *P2* mRNA levels in M1 and M2 macrophages by percentage and mean fluorescence intensity (MFI) in decidua. Results based on 5 uninfected dams. Student’s T-test used to calculate p-values.

## Discussion

A unique challenge in both the operation as well as the study of immune responses occurring during pregnancy is that protection for the host may not always encompass the fetus. While the placenta acts as an immunological and physical barrier for fetal protection, it is susceptible to pathogens capable of colonizing this tissue and, under worsening conditions, can reorient towards an exclusively maternal protective response (Robbins and Bakardjiev 2012; Zeldovich and Bakardjiev 2012; Bonney and Johnson 2019; Jena et al. 2019). Decidual macrophages have been shown to be critical for the plasticity of the placental response in being important for both the maintenance of fetal tolerance as well as responding to inflammatory and/or infectious conditions that can result in immune activation against the fetus (Brown et al. 2014; Wang et al. 2018).

We show here that Perforin-2 plays a significant role during *L. monocytogenes* infection in the placenta and fetus. We additionally found that fetal-derived Perforin-2 in the placenta contributes to limiting *L. monocytogenes* infection, although there is a dominance in regards to the maternal expression of Perforin-2. These findings are consistent with published data showing that the maternal-derived decidua is the initial site of infection and the first line of defense in bloodborne *L. monocytogenes* infections (Rizzuto et al. 2017). In analysis of *Perforin-2* mRNA levels in individual decidual cells, we found that CD45^+^ cells possessed abundant transcripts, and specific to decidual macrophages, these levels became elevated in infections with high pathogen loads. This inductive *Perforin-2* expression in decidual macrophages correlated with their polarization from the tolerogenic M2 phenotype to the inflammatory M1 phenotype. In these experiments with high pathogen loads (i.e., PD_100_; 5 – 10 × 10^5^ CFU), dams often displayed typical signs of listeriosis at 44 hpi including ruffled fur, hunched backs, and slowed movement. Also there would occasionally be FPU observed in the cage housing presenting with darkened uterine horns and loss of fetal structures in remaining fetuses, all indicative of maternal immune activation (MIA) (Goldstein et al. 2017). In contrast, there was little to no inductive *Perforin-2* expression in decidual macrophages in infections with relatively lower pathogen loads (i.e., PD_50_; 2 × 10^5^ CFU) in which neither disease symptoms nor fetal expulsions occur. These findings indicate that inductive *Perforin-2* expression in deciduae is associated with a pathological shift in macrophage phenotype.

In understanding *Perforin-2* expression in decidual macrophages, it is worth noting that there are three immunological stages of pregnancy (Mor et al. 2017). At the early stage of pregnancy, when the blastocyst is implanted into the uterine wall, there is increased inflammation and a Th_1_-biased environment, during which there is a predominance of M1 macrophages in the decidua. It has been shown that this inflammation is not in response to the invading fetal antigens but that there is an active recruitment by the trophoblast cells in an effort to educate immune cells towards fetal tolerance. The mid-stage of pregnancy is a period of fetal growth and when fetal tolerance is the most important, as pathological conditions that lead to MIA can result in a breach of fetal tolerance and therefore fetal rejection (Mor et al. 2017). In line with this, the mid-term of pregnancy represents an anti-inflammatory state, generating a tolerogenic Th_2_-type immune environment in which M2 macrophages dominate within deciduae. Lastly, the late stage of pregnancy again requires inflammation that is necessary for labor induction and is characterized by a Th_1_ immune response in which again there is a predominance of M1 macrophages.

Pathogens capable of crossing the placental barrier can take advantage of the anti-inflammatory microenvironment of the placenta during the mid-stage of pregnancy. However, once a certain threshold is reached and MIA occurs, fetal tolerance can be disrupted and result in pregnancy termination (Bonney and Johnson 2019). Perforin-2 serves both a protective function in sub-MIA infections (i.e., as shown in the PD_50_ experiments [Figs. 1 and 2]) as well as becoming elevated during infections in which MIA is triggered (Fig. 5). Collectively, our investigation may indicate that Perforin-2 is a component that acts to protect either both the mother and fetus (sub-MIA infections) or exclusively the mother (MIA infections).

